# A critical role of brain network architecture in a continuum model of autism spectrum disorders spanning from healthy individuals with genetic liability to individuals with ASD

**DOI:** 10.1101/2021.09.03.458876

**Authors:** Budhachandra Khundrakpam, Neha Bhutani, Uku Vainik, Noor Al-Sharif, Alain Dagher, Tonya White, Alan C. Evans

**Affiliations:** McGill Centre for Integrative Neuroscience, Montreal Neurological Institute, McGill University, Montreal, QC, Canada H3A 2B4; Montreal Neurological Institute, McGill University, Montreal, QC, Canada H3A 2B4; Institute of Psychology, Faculty of Social Sciences, University of Tartu, Estonia, Naituse 2-216, 50409; Erasmus MC-Sophia / Kamer KP-2869, Postbus 2060, 3000 CB Rotterdam

## Abstract

Studies have shown cortical alterations in individuals with autism spectrum disorders (ASD) as well as in individuals with high polygenic risk for ASD. An important addition to the study of altered cortical anatomy is the investigation of the underlying brain network architecture that may reveal brain-wide mechanisms in ASD and in polygenic risk for ASD. Such an approach has been proven useful in other psychiatric disorders by revealing that brain network architecture shapes (to an extent) the disorder-related cortical alterations. This study uses data from a clinical dataset – 560 male subjects (266 individuals with ASD and 294 healthy individuals, CTL, mean age at 17.2 years) from the Autism Brain Imaging Data Exchange database, and data of 391 healthy individuals (207 males, mean age at 12.1 years) from the Pediatric Imaging, Neurocognition and Genetics database. ASD-related cortical alterations (group difference, ASD-CTL, in cortical thickness) and cortical correlates of polygenic risk for ASD were assessed, and then statistically compared with structural connectome-based network measures (such as hubs) using spin permutation tests. Next, we investigated whether polygenic risk for ASD could be predicted by network architecture by building machine-learning based prediction models, and whether the top predictors of the model were identified as disease epicenters of ASD. We observed that ASD-related cortical alterations as well as cortical correlates of polygenic risk for ASD implicated cortical hubs more strongly than non-hub regions. We also observed that age progression of ASD-related cortical alterations and cortical correlates of polygenic risk for ASD implicated cortical hubs more strongly than non-hub regions. Further investigation revealed that structural connectomes predicted polygenic risk for ASD (*r*=0.30, *p*<0.0001), and two brain regions (the left inferior parietal and left suparmarginal) with top predictive connections were identified as disease epicenters of ASD. Our study highlights a critical role of network architecture in a continuum model of ASD spanning from healthy individuals with genetic risk to individuals with ASD. Our study also highlights the strength of investigating polygenic risk scores in addition to multi-modal neuroimaging measures to better understand the interplay between genetic risk and brain alterations associated with ASD.

## Introduction

Neuroimaging studies have consistently shown alterations in cortical anatomy in individuals with autism spectrum disorders (ASD) (Hadjikhani et al. 2006; Hyde et al. 2010; Raznahan et al. 2010; Wallace et al. 2010; Ecker et al. 2013; Zielinski et al. 2014; Lange et al. 2015; Khundrakpam, Lewis, Kostopoulos, et al. 2017; Bezgin et al. 2018; Van Rooij et al. 2018). For example, a recent large-scale study out of the ENIGMA consortium (3222 individuals, 1571 with AUTISM) showed increased cortical thickness in the frontal cortex (Cohen’s *d* = 0.20) and decreased thickness in the temporal cortex (Cohen’s *d* = −0.21) in individuals with ASD (Van Rooij et al. 2018). Interestingly, the ASD-related cortical alterations extend beyond those with clinical diagnosis to the general population (Di Martino et al. 2009; Barnea-Goraly et al. 2010; Blanken et al. 2015, 2017), consistent with an emerging framework that conceptualizes ASD as a continuum model (with a normal distribution of autistic tendencies in the general population where a full diagnosis is at the severe tail of the distribution (Constantino and Todd 2003; Wakabayashi et al. 2006; Plomin et al. 2009; Constantino 2011; Robinson et al. 2011, 2016; Blanken et al. 2015, 2017; Hyseni et al. 2019)). For example, cortical thickness of the frontal and parietal regions were dimensionally related to genetic risk for ASD in general population, and were also part of the cortical alterations associated with ASD in clinical population (Khundrakpam et al. 2020).

An important addition to the study of altered cortical anatomy in ASD is the investigation of the underlying brain network architecture that may reveal brain-wide mechanisms in ASD. Such an approach has been proven useful in other psychiatric disorders by revealing that brain network architecture shapes (to an extent) the disorder-related morphological alterations (Zhou et al. 2012; Larivière, Rodríguez-cruces, et al. 2020). Of particular interest is the study of brain *hubs* (regions with several connections that serve as information relay centers (Bullmore and Sporns 2009; van den Heuvel and Sporns 2013)) which have been implicated in several psychiatric disorders (Crossley et al. 2014; Baker et al. 2015; Fornito et al. 2015; Larivière, Rodríguez-cruces, et al. 2020). These studies revealed that cortical alterations associated with the disorders were greater in the hubs compared to the peripheral regions (with only local connections), possibly due to the high metabolic activity and their links with several brain networks (Buckner et al. 2009; Zhou et al. 2012; Crossley et al. 2014). In addition to examining hubs, recent studies have also identified *disease epicenters*, defined as regions whose network architecture play central role in the whole-brain manifestation of psychiatric disorders (Zhou et al. 2012; Zeighami et al. 2015; Filippi et al. 2020; Larivière, Rodríguez-Cruces, et al. 2020; Shafiei et al. 2020). Taken together, investigation of hubs and disease epicenters may provide novel insights into how patterns of ASD-related cortical alterations may be configured by the brain network architecture.

We, therefore, set out to examine the relation between brain network architecture and ASD-related alterations in cortical anatomy. Extending this goal in the context of the continuum model, we also set out to examine the link between brain network architecture and cortical correlates of polygenic risk for ASD. For this, *i)* ASD-related cortical alterations (group difference, ASD-CTL in cortical thickness) using a clinical cohort and *ii*) cortical correlates of polygenic risk for ASD using a general population, were first computed. Next, using normative structural brain networks derived from diffusion magnetic resonance imaging (dMRI) data from general population, we then tested the hypothesis – whether there was a selective vulnerability of hub regions that parallel the ASD-related cortical alterations as well as the cortical correlates of polygenic risk for ASD. Lastly, we set out to investigate whether polygenic risk for ASD could be predicted by structural brain networks, and whether the top predictors of the model corresponded with the disease epicenters of ASD.

## Methods

### Subjects

Data for the study were taken from two publicly available databases: *i*) a clinical cohort: the Autism Brain Imaging Data Exchange (ABIDE) database (Di Martino et al. 2014), and *ii*) a general population cohort: the Pediatric Imaging, Neurocognition and Genetics (PING) study (Jernigan et al. 2016). While ABIDE is an agglomerative dataset of MRI scans of healthy individuals and individuals with ASD (Di Martino et al. 2014), the PING study comprises of neuroimaging, cognition and genetic data from 1493 typically developing children and adolescents collected from 10 different sites across the United States (Jernigan et al. 2016).

### Genomic data and computation of polygenic risk scores

The PING dataset includes 550,000 single nucleotide polymorphisms (SNPs) genotyped from saliva samples using Illumina Human660W-Quad BeadChip. Computation of polygenic risk scores (PRS) followed steps similar to that of our previous study (Khundrakpam et al. 2020). Summary of steps include: preparation of the data for imputation using the “imputePrepSanger” pipeline (https://hub.docker.com/r/eauforest/imputeprepsanger/) and implemented on CBRAIN (Sherif et al. 2014) using Human660W-Quad_v1_A-b37-strand chip as reference. The next step involved data imputation with Sanger Imputation Service (McCarthy et al. 2016), using default settings and the Haplotype Reference Consortium, HRC (http://www.haplotype-reference-consortium.org/) as the reference panel. Using Plink 1.9 (Chang et al. 2015), the imputed SNPs were then filtered with the inclusion criteria: *i*) SNPs with unique names, *ii*) only ACTG and *iii*) MAF > 0.05. All SNPs that were included had INFO scores R^2^ > 0.9 with Plink 2.0. Next, using polygenic score software PRSice 2.1.2 (Euesden et al. 2015) additional ambiguous variants were excluded, resulting in 4,696,385 variants being available for polygenic scoring. We filtered individuals with 0.95 loadings to the European principal component (GAF_Europe variable provided with the PING data), resulting in 526 participants. These participants were then used to compute 20 principal components with Plink 1.9. The polygenic risk score for ASD was based on ASD GWAS trained on 18,381 <u>independent</u> individuals with ASD and 27,969 controls (Grove et al. 2019). Similar to our previous study (Khundrakpam et al. 2020), the data was clumped based on PRSice default settings (clumping distance = 250kb, threshold r2 = 0.1), using *p* = 0.001 cut-off criterion. After matching with available variants in the data, the polygenic risk score for ASD was based on 1245 variants.

### Image pre-processing and quality control

For structural MRI data (of both the ABIDE and PING datasets), we used the CIVET processing pipeline, (https://mcin.ca/technology/civet/) developed at the Montreal Neurological Institute to compute cortical thickness measurements at 81,924 regions covering the entire cortex. Summary of steps include; non-uniformity correction of the *T_1_*-weighted image and then linear registration to the Talairach-like MNI152 template (created with the ICBM152 dataset). After repeating the non-uniformity correction using template mask, the non-linear registration from the resultant volume to the MNI152 template is computed, and the transform used to provide priors to segment the image into GM, WM, and cerebrospinal fluid. Inner and outer GM surfaces are then extracted using the Constrained Laplacian-based Automated Segmentation with Proximities (CLASP) algorithm. Cortical thickness is then measured in native space using the linked distance between the two surfaces at 81,924 vertices. In order to impose a normal distribution on the corticometric data and to increase the signal to noise ratio, each individual’s cortical thickness map was blurred using a 30-millimeter full width at half maximum surface-based diffusion smoothing kernel. Two independent reviewers performed quality control (QC) of the data, and only scans with consensus of the two reviewers were used. Exclusion criteria for QC procedure include - data with motion artifacts, a low signal to noise ratio, artifacts due to hyperintensities from blood vessels, surface-surface intersections, or poor placement of the grey or white matter (GM and WM) surface for any reason.

For diffusion MRI data (of the PING dataset), we used the FSL pipeline (FMRIB Software Library v5.0.9) (Jenkinson et al. 2012) for pre-processing. Summary of steps include; correction of the distortion effects induced by eddy currents, inter-volume movements and susceptibility of the diffusion data; rigid alignment of the individual unweighted image with the structural image using flirt; non-linear registration to transform individual structural image to an MNI152 standard T_1_-weighted template using fnirt; computing the forward and backward warp field images between individual dMRI and MNI T_1_ spaces by concatenating (or inverting) the rigid transformation matrix and the warp field image. Diffusion parameters at each voxel were estimated by using Markov Chain Monte Carlo sampling. In this step, up to 2 possible fiber populations were modeled for each voxel after 2000 iterations. Quality control of the data was performed by checking the structural image and the average of the non-diffusion-weighted images for each individual, and then evaluation of the results of registration by visual inspection. Exclusion criteria of QC include – if signal-noise-rate (SNR) of structural image or unweighted-diffusion image was lower than 800, and data with greater than 2 mm frame-wise displacements of the dMRI.

### Sample characteristics

For the clinical cohort (ABIDE dataset), apart from the QC procedure, there were additional exclusion criteria namely insufficient number of individuals in each diagnostic group (ASD and CTL) to determine group difference, too few females resulting to excluding all females, and excluding individuals over 35 years of age due to insufficient numbers. The final sample consisted of 560 male individuals, 266 individuals with ASD (17.2±6.4 years) and 294 controls (17±6.4 years) (**Table 1A**).

**Table 1:** Description of study sample. A) Clinical sample from the ABIDE dataset, B) General population sample from the PING dataset. Note, age is given as mean±std, ABIDE = Autism Brain Imaging Data Exchange, PING = Pediatric Imaging, Neurocognition and Genetics, PRS = Polygenic risk score, FSIQ = full-scale intelligence quotient, NTCB = The NIH Toolbox Cognition Battery.

| **A. ABIDE sample** |  | **B. PING sample** |  |
| --- | --- | --- | --- |
| Number of subjects, N | 560 | Number of subjects, N | 391 |
| Males | 560 | Males/females | 207/184 |
| ASD/CTL | 266/294 | Age (in years) | 12.1 ± 4.7 |
| Age-ASD (in years) | 17.2 ± 6.4 | PRS | 5.5e-03 ± 5.8e-04 |
| Age-CTL (in years) | 17.0 ± 6.4 | NTCB Reading score | 134.4 ± 70.6 |
| FSIQ-ASD | 106.4 ± 15.8 | NTCB Flanker score | 7.8 ± 1.7 |
| FSIQ-CTL | 111.4 ± 12.1 | NTCB DCCS score | 7.8 ± 1.3 |

For the general population (PING dataset), of the total 1493 individuals, filtering for individuals with 0.95 loadings to the European principal component resulted in 526 individuals. Of these, 95 individuals did not have MRI data and 2 subjects did not have information about age, resulting in 429 subjects. Next, 13 subjects were excluded before any processing (raw data) due to severe motion and slicing artifacts. A subsequent 25 subjects failed CIVET pipeline (for a number of reasons including presence of bright blood vessels and poor contrast) and were excluded in further analysis. The final sample consisted of 391 participants (males/females=207/184, age=12.1±4.7 years) (**Table 1B**).

### Determination of hubs

The structural connectivity matrices were used to determine hubs (regions with several connections that serve as information relay centers) by computing centrality maps based on graph-theoretic measures (e.g. degree centrality) (Sporns et al. 2007). Computations were done using the Brain Connectivity Toolbox (https://sites.google.com/site/bctnet/).

### Statistical analysis of regional overlap

Since hub regions have been shown to be more susceptible to disorder-related cortical alterations than non-hub regions, we next investigated whether the observed cortical differences were influenced by the network connectivity of brain regions. For this, we statistically checked regional overlap between *i*) map of the group difference (ASD-CTL) in cortical thickness (using the ABIDE dataset) and *ii*) centrality map. In a similar manner, we statistically checked regional overlap between *i*) map of the association of polygenic risk for ASD and cortical thickness (using the PING dataset) and *ii*) centrality map. The statistical comparisons were done using the spin test developed by (Alexander-Bloch et al. 2018). In short, the method, using a spatial permutation framework, generates null models of overlap by applying random rotations to spherical representations of the cortical surface. As in previous studies (Reardon et al. 2018), 1000 surface rotations of the PING map were generated, and the statistical overlap was checked by comparing whether the observed cross-vertex correlation between two maps was statistically greater (*p*<0.05) than those with 1000 rotations. Similar analysis was done for statistically comparing the centrality maps and *i*) map of age*(ASD-CTL) on cortical thickness (using the ABIDE dataset) and *ii*) map of age*(PRS) on cortical thickness (using the PING dataset).

### Connectome predictive modeling of polygenic risk for ASD

With structural connectivity data as input, we used the connectome predictive modeling (CPM) approach to investigate whether structural connectomes can be used to predict PRS-ASD. Recently introduced, CPM approach (Finn et al. 2015; Shen et al. 2017) is a data-driven framework which utilizes cross-validation to build predictive models of brain-behaviour associations from connectivity data. Comprising of four steps – feature selection, feature summarization, model building and computation of prediction significance (Finn et al. 2015; Shen et al. 2017), the framework has been validated and used in predicting anxiety (Wang et al. 2020), attention (Yoo et al. 2018), maternal bonding (Rutherford et al. 2020), etc. The accuracy of the prediction model was assessed by computing correlation between the true and predicted PRS-ASD scores. Next, the top predictors of the model were visualized using the BrainNet Viewer (Xia et al. 2013).

### Computation of disease epicenters of ASD

In order to check whether the top predictors of the model were identified as disease epicenters of ASD, we first compared each brain region’s structural connectivity with the ASD-related cortical alterations. Next, we used spin permutation tests to compute the significance of the correlations. Brain regions with statistically significant correlations were identified as disease epicenters of ASD. Lastly, we checked whether any of the identified disease epicenters of ASD corresponded with the top predictors of PRS-ASD.

## Results

### Hub organization is associated with ASD-related cortical alterations as well as cortical correlates of polygenic risk for ASD

We asked whether network organization was associated with ASD-related cortical alterations (group difference in cortical thickness between ASD and CTL). For this, we used structural connectivity data (derived from diffusion-weighted tractography) from the PING dataset. Using structural connectome data, hubs (brain regions with larger degree centrality) were identified. As in earlier studies of network centrality maps in healthy individuals (Zuo et al. 2012; van den Heuvel and Sporns 2013), hubs were localized in the medial prefrontal, superior parietal and superior temporal regions (**Figure 1A**). ASD-related cortical difference maps were obtained by computing the group difference (ASD-CTL) in cortical thickness using the ABIDE dataset (for details, see **Methods**). Consistent with previous studies (Khundrakpam, Lewis, Kostopoulos, et al. 2017), greater cortical thickness was observed in several brain regions including the left parietal and lateral frontal and bilateral temporal regions in individuals with ASD (**Figure 1A**). Analysis of spatial similarity between ASD-related cortical difference pattern and degree centrality map was compared through correlation analysis (and statistically assessed via non-parametric spin permutation tests, see **Methods**), and revealed that ASD-related cortical difference implicated cortical hubs (*r* = 0.31, *p*_spin_ = 0.015) more strongly than non-hub regions (**Figure 1A**). Cortical correlates of polygenic risk for ASD (association of cortical thickness and PRS for ASD) also implicated cortical hubs (*r* = 0.37, *p*_spin_ = 0.003) more strongly than non-hub regions (**Figure 1B**).

**Figure 1.**
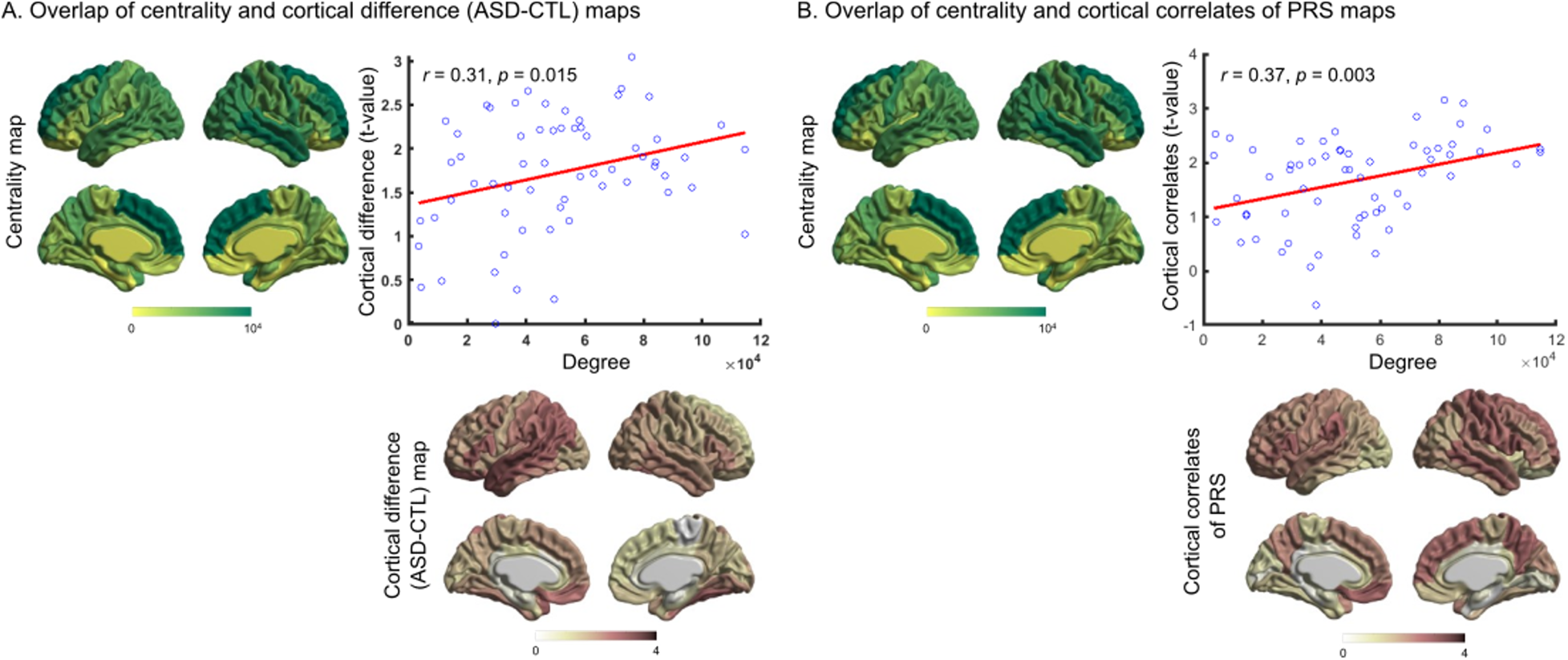
Association of structural hub organization and ASD-related cortical alterations and cortical correlates of polygenic risk for ASD. (A) Significant correlation (*r*_spin_=0.31, *p*=0.015) was observed between centrality and ASD-related cortical difference maps (group difference, ASD-CTL, in cortical thickness). (B) Significant correlation (*r*_spin_=0.37, *p*=0.003) was observed between centrality and cortical correlates of polygenic risk for ASD (association of cortical thickness and polygenic risk for ASD) maps. Note, ASD = autism spectrum disorders, CTL = controls, PRS = polygenic risk score.

### Hub organization is associated with cross-sectional progression of ASD-related cortical alterations as well as cortical correlates of polygenic risk for ASD

The interaction of age and difference in cortical thickness for individuals with ASD compared to controls can give insights into the cross-sectional progression of ASD-related cortical alterations. By extension, the interaction of age and cortical correlates of polygenic risk for ASD could indicate cross-sectional progression of polygenic risk-related cortical correlates. Thus, using general linear models, we first examined effect of age on *i*) ASD-related cortical difference map and *ii*) cortical correlates of polygenic risk for ASD. Comparison of the age interaction patterns of ASD-related cortical alterations and degree centrality maps showed significant correlations with the cortical hubs (*r* = 0.38, *p*_spin_ = 0.002, **Figure 2A**). Similarly, comparison of the age interaction patterns of cortical correlates of polygenic risk for ASD and degree centrality maps showed significant correlations with the cortical hubs (*r*= 0.43, *p*_spin_ = 0.0005, **Figure 2B**).

**Figure 2.**
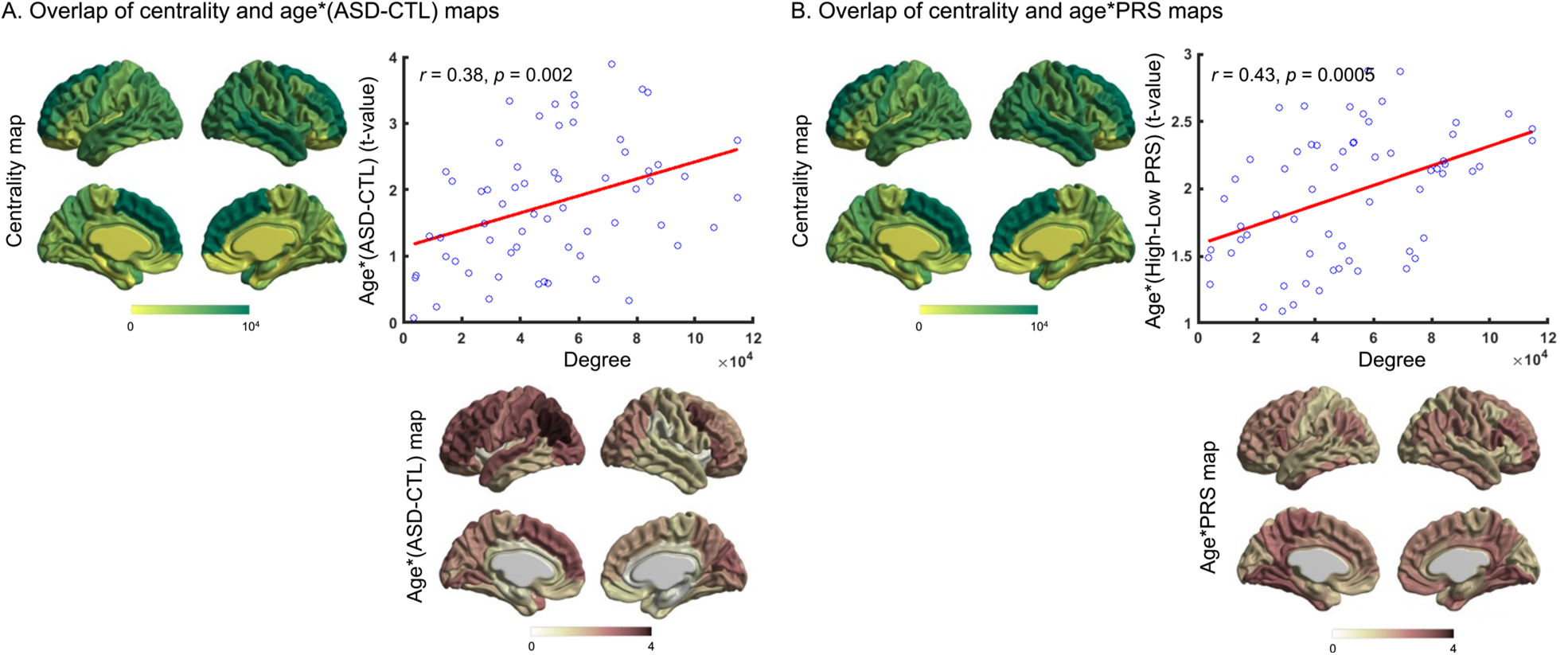
Association of structural hub organization and cross-sectional progression of ASD-related cortical alterations and cortical correlates of polygenic risk for ASD. (A) Significant correlation (*r*_spin_=0.38, *p*=0.002) was observed between age interaction patterns of ASD-related cortical alterations and degree centrality maps. (B) Significant correlation (*r*_spin_=0.43, *p*=0.0005) was observed between age interaction patterns of cortical correlates of polygenic risk for ASD and degree centrality maps. Note, ASD = autism spectrum disorders, CTL = controls, PRS = polygenic risk score.

### Connectome predictors of polygenic risk for ASD relate to disease epicenters of ASD

Using the Connectome Predictive Modeling (CPM) framework, structural connectivity data predicted polygenic risk for ASD with *r* = 0.30, *p* < 0.0001 (**Figure 3A**) suggesting that ~ 9% of the variance in PRS-ASD may be explained by structural connectivity. Further investigation revealed the most predictive features (brain connections) comprising of ipsilateral connections (the left frontal-frontal, parietal-parietal connections and right temporal-temporal connections) and bilateral postcentral connections (**Table 2, Figure 3C**). We next checked whether any of these regions (with top predictive connections) were identified as epicenters of ASD. For this, as outlined in **Methods**, disease epicenters for ASD were computed (**Figure 3D**). We found that two of the identified epicenters (left inferior parietal and left suparmarginal) corresponded with regions with top predictive connections (**Figure 3D**). In these two regions, ASD-related cortical alterations were significantly associated with seed-based structural connectivity of the regions (**Figure 3B**).

**Figure 3.**
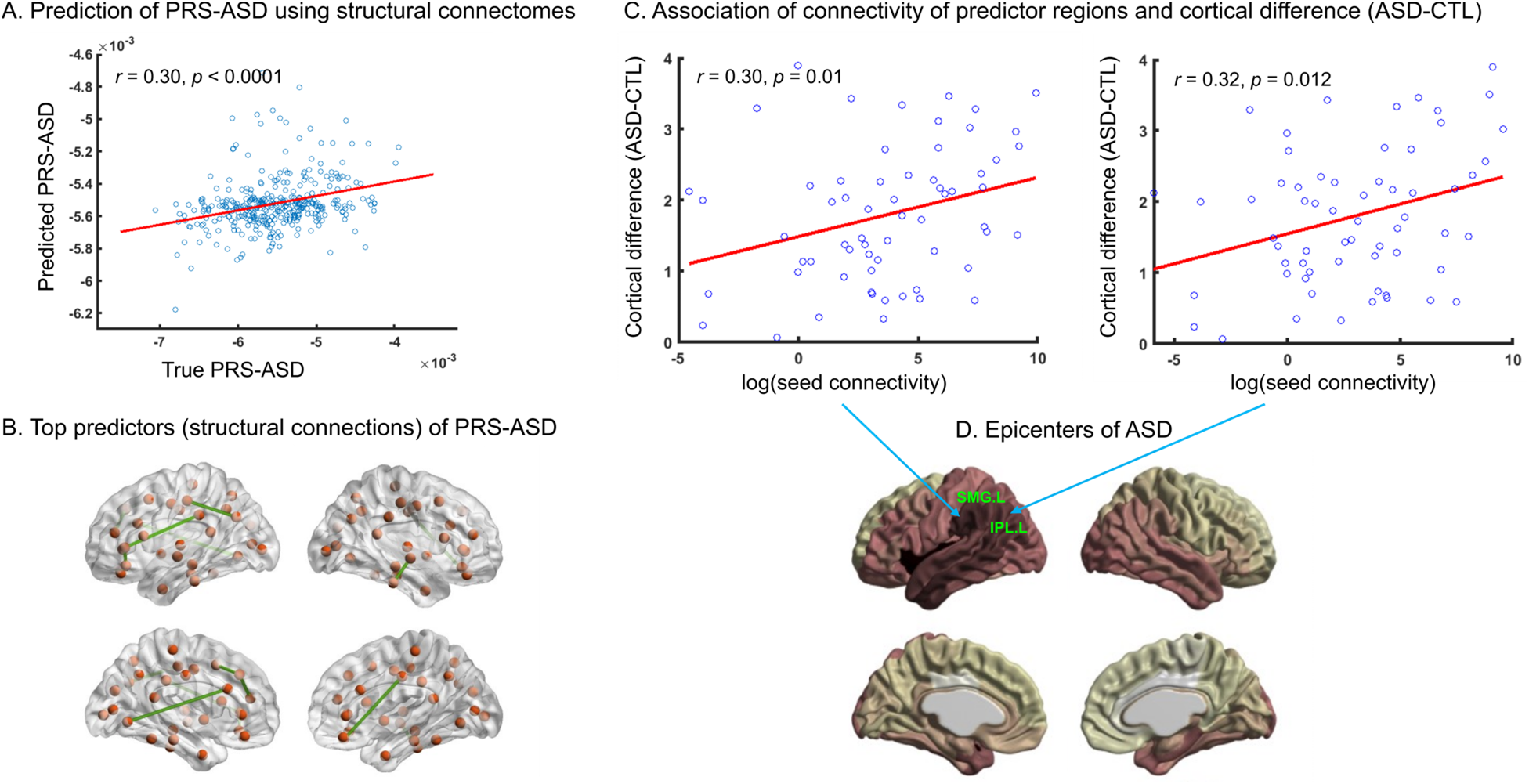
Relation of connectome predictors of PRS-ASD and disease epicenters of ASD. (A) Connectome predictive modeling using structural connectivity resulted in accurate prediction of PRS-ASD (*r*=0.30, *p*<0.0001 between true and predicted PRS-ASD). (B) Top predictors of the model included ipsilateral connections (the left frontal-frontal, parietal-parietal connections and right temporal-temporal connections) and bilateral postcentral connections. (C-D) Of these top predictors, two regions – SMG.L and IPL.L were found to be epicenters of ASD where connectivity of the region were significantly related to ASD-related cortical alterations (see **Methods**). Note, ASD = autism spectrum disorders, CTL = controls, PRS = polygenic risk score, SMG.L = left supramarginal gyrus, IPL.L = left inferior parietal lobule.

**Table 2:** Brain connections which were identified as top predictors of PRS-ASD. Note, ASD = autism spectrum disorders, PRS = polygenic risk score.

| Region | Region |
| --- | --- |
| Left Caudal Anterior Cingulate | Left Lingual |
| Left Pars Orbitalis | Left Pars Triangularis |
| Left Inferior Parietal | Left Postcentral |
| Left Caudal Middle Frontal | Left Superior Frontal |
| Left Rostral Middle Frontal | Left Superior Frontal |
| Left Pars Triangularis | Left Supramarginal |
| Left Postcentral | Right Postcentral |
| Right Lateral Orbito-Frontal | Right Posterior Cingulate |
| Right Inferior Temporal | Right Superior Temporal |

## Discussion

In this study, using large-scale MRI data from a clinical cohort and a general population sample, we examined whether there were links between brain network architecture and ASD-related cortical alterations. Additionally, on the basis of the context of the continuum model, we also investigated whether brain network architecture also has links with cortical correlates of polygenic risk for ASD. We observed that ASD-related cortical alterations implicated cortical hubs more strongly than non-hub regions. Similarly, cortical correlates of polygenic risk for ASD also implicated cortical hubs more strongly than non-hub regions. Comparison of the age interaction patterns of ASD-related cortical alterations with degree centrality maps showed significant correlations with the cortical hubs. Similarly, comparison of the age interaction patterns of cortical correlates of polygenic risk for ASD with degree centrality maps showed significant correlations with the cortical hubs. Further investigation revealed that structural connectivity predicted polygenic risk for ASD (*r* = 0.30, *p* < 0.0001) suggesting that ~ 9% of the variance in polygenic risk for ASD may be explained by structural connectivity. Further investigation revealed two of the disease epicenters (the left inferior parietal and left suparmarginal) as regions with the top predictive connections. Taken together, our findings suggest a critical role of network architecture in ASD-related cortical alterations and in cortical correlates related to polygenic risk for ASD.

Several studies have consistently shown cortical alterations in individuals with ASD (Hadjikhani et al. 2006; Hyde et al. 2010; Raznahan et al. 2010; Wallace et al. 2010; Ecker et al. 2013; Zielinski et al. 2014; Lange et al. 2015; Khundrakpam, Lewis, Kostopoulos, et al. 2017; Bezgin et al. 2018; Van Rooij et al. 2018). Our findings add to the extant literature of ASD research by revealing how brain network architecture is linked with the cortical alterations associated with ASD. Firstly, we observed that regions with more hub characteristics (greater centrality) showed greater ASD-related cortical alterations, in consistent with previous studies that have implicated hubs in brain disorders (Crossley et al. 2014; Baker et al. 2015; Fornito et al. 2015; Larivière, Rodríguez-cruces, et al. 2020). Secondly, in consistent with previous studies (Zeighami et al. 2015; Larivière, Rodríguez-cruces, et al. 2020), we observed that brain regions with the largest ASD-related cortical alterations denote disease epicenters for ASD (**Fig. 3C**). Thirdly, we observed that brain regions with high degree centrality exhibited greater cross-sectional progression of ASD-related cortical alterations (**Fig. 2D**).

The epicenters identified in our study encompass somatosensory, motor and visual areas along with regions involved in auditory processing and low-level sensory integration (**Fig. 3D**). Individuals with ASD usually show impairments in motor behaviors, responses to tactile, auditory, and visual stimuli, and in their processing of language and nonlinguistic social stimuli (Tager-Flusberg 1981; Schultz et al. 2003; Schultz 2005; Leekam et al. 2007; Redcay 2008). More interestingly, the two epicenters (suparmarginal gyrus and inferior parietal lobule) identified as top predictors of polygenic risk for ASD are involved in visuospatial processing. These brain regions, located at the intersection of the visual, auditory and somatosensory cortices, comprise of neurons with multimodal properties which can process several stimuli concurrently. Inferior parietal lobule is involved in integration and interpretation of sensory information, emotional perception of sensory stimuli. The inferior parietal lobule, with its connections to both Broca’s area and Wernicke’s area, may serve as information relay center between these areas for language-related functions. The inferior parietal lobule, part of the social brain system, is also involved in social perception (Pelphrey and Carter 2008) and executive attention (Corbetta et al. 2008). Not surprisingly, alterations in the connectivity of the inferior parietal lobule are linked with deficits in social cognition in ASD (Hadjikhani et al. 2006; Cheng et al. 2011). Lastly, altered white matter in the inferior parietal lobule has been shown in children with ASD (Yang et al. 2018) which in turn has been associated with impaired motor performance (Hanaie et al. 2016).

The mechanisms behind the influence of brain network architecture on ASD-related cortical alterations are not clear. In terms of cortical alterations, several factors including larger neurons, increased number of neurons (Courchesne et al. 2011), greater microglial cell density and somal volume (Morgan et al. 2010) and greater number of synaptic spines and reduced developmental synaptic pruning (Tang et al. 2014) have been associated with increased cortical thickness in individuals with ASD. On the other hand, alterations in white matter connectivity in ASD might be due to increased packing density (Bauman and Kemper 2005), increased oedema from inflammation (Vargas et al. 2005) and reduced thickness of myelin (Zikopoulos and Barbas 2010). Additionally, reduced synaptic pruning seen in children with ASD might hinder axonal remodeling resulting in altered connectivity (Auerbach et al. 2011; Tang et al. 2014). Further, abnormalities in social perception and executive attention (which are considered broad phenotypes for ASD) might impact processing of neural information, which in turn might alter brain structure and connectivity (Valla and Belmonte 2013).

Our findings that network architecture is linked to ASD-related cortical alterations in clinical cohort as well as cortical correlates of ASD polygenic risk in general population highlight an emerging consensus that psychiatric disorders including ASD may be viewed as continuum models as opposed to conventional diagnostic groups. Traditionally, psychiatric disorders are categorized as diagnostic groups: *individuals with disorder (case)* and *healthy individuals (control)*. However, recent studies have indicated that psychiatric disorders may be viewed as a continuum with a normal distribution of psychiatric tendencies in the general population, where a full diagnosis is at the severe tail of the distribution (Constantino and Todd 2003; Wakabayashi et al. 2006; Plomin et al. 2009; Constantino 2011; Robinson et al. 2011, 2016; Blanken et al. 2015, 2017; Hyseni et al. 2019). Evidence toward this has come from behavioural and imaging studies. For instance, autistic traits (e.g. social and communications deficits) extend beyond diagnostic groups into the general population (Constantino and Todd 2003). In terms of imaging, brain alterations have been observed not only for case-control differences (Raznahan et al. 2010; Ecker et al. 2013; Lange et al. 2015; Khundrakpam, Lewis, Reid, et al. 2017; Van Rooij et al. 2018), but also for autistic traits in the general population (Di Martino et al. 2009; Barnea-Goraly et al. 2010; Blanken et al. 2015, 2017). One such study found significant negative association of cortical thickness (in the right superior temporal cortex) and a continuous measure of autistic traits in a large longitudinal sample of normally developing youths (Wallace et al. 2012). In another study, Blanken et al. observed significant association of gyrification and autistic traits along a continuum in a large populationbased sample of children (Blanken et al. 2015). Recently, we found that cortical correlates of polygenic risk for ASD in general population overlap with the cortical alterations seen in individuals with ASD (Khundrakpam et al. 2020). Extending these previous studies, our findings indicate that brain network architecture is linked to ASD-related cortical alterations in clinical population as well as cortical correlates of ASD polygenic risk in general population. Our observation that structural connectivity could accurately predict polygenic risk for ASD and more importantly, that two of the top predictor regions were identified as disease epicenters of ASD lends more credence on the continuum model of ASD spanning from healthy individuals with genetic risk to individuals with ASD.

In conclusion, our study highlights a critical role of network architecture in ASD-related cortical alterations and in cortical correlates of polygenic risk for ASD. Our study also underscores the need for investigating polygenic risk scores in addition to multi-modal neuroimaging measures to better understand the interplay between genetic risk and brain abnormalities associated with ASD.

## Acknowledgements

BK was supported by a Brain & Behavior Research Foundation (BBRF) – NARSAD 2020 YI Grant (Grant Number 29492). ACE was supported by a CIHR Grant (Fund 254573). UV has been funded by Estonian Research Council’s personal research funding start-up grant PSG656.

